# Not all interactions are positive: testing how antagonistic interactions impact the robustness of plant-pollinator networks

**DOI:** 10.1101/538702

**Authors:** Heather M. Briggs, Carolyn A. Ayers, Paul R. Armsworth, Berry J. Brosi

## Abstract

Given ongoing pollinator declines, it is important to understand the dynamics of linked extinctions of plants driven by pollinator extinctions. Topological robustness models focused on this question suggest relatively high robustness of plant species to pollinator species extinctions. Still, existing robustness models typically assume that all interactions in plant-pollinator networks are positive, which is clearly not always the case. For example, many pollinators remove floral resources without transferring pollen, or even damage floral structures in the case of nectar robbing. Here we introduce antagonistic interactions into plant-pollinator networks and assess the resilience of plant communities to pollinator species losses. Incorporating antagonistic interactions leads to lower network robustness, i.e. an increased rate of plant species loss (as compared to networks with only mutualistic interactions) in empirical plant-pollinator networks. In conjunction with extinction order, the addition of increasingly antagonistic interactions was idiosyncratic in that it did not always magnify the effects of extinction order across the three networks. These results underscore the importance of considering the full spectrum of interaction outcomes when assessing robustness to coextinctions in plant-pollinator networks, as well as other ecological systems.

## Introduction

Pollinator losses are increasing across the globe which could have potentially strong negative effects on the plants that rely on them for pollination (Biesmeijer et al. 2006, Potts et al. 2010). Network-based simulations suggest that plant communities will be very robust to pollinator extinctions (i.e. secondary plant extinctions resulting from pollinator extinctions) (e.g. Memmott et al. 2004, 2007, Kaiser-Bunbury et al. 2010). This robustness is likely driven by two features of network structure. First, pollination networks are dominated by *generalist* interactions; most plants and pollinators each interact with several species from the other group over the course of their lives (Waser et al. 1996). Second, pollination networks have a *nested* structure in which specialist plants and pollinators tend to interact with a subset of the species in the other group that generalists interact with (Bascompte et al. 2003, 2006). Nestedness leads to an asymmetric interaction structure where specialists from one group tend to act with generalists, not specialists, from the other, which could reduce linked extinctions if specialists are vulnerable to stochastic extinctions (Bascompte et al. 2006, Bascompte & Jordano 2007). One feature of binary-graph plant-pollinator simulation models that may overestimate network robustness is that all interactions in a binary-graph network are positive (i.e. strictly mutualistic). A typical binary-graph simulation modeling approach assumes that if at least one link remains between a plant and a pollinator, the plant will continue to persist (Memmott et al. 2004, 2007).

While the assumption of “all interactions positive” is a reasonable starting point for mutualistic network models, empirical evidence suggests at least three ways in which pollinators can have negative consequences for the reproduction of plants they interact with (Irwin et al. 2010, Ashman & Arceo-Gomez 2013, Brosi & Briggs 2013). First, some pollinators may visit and extract nectar or pollen rewards from flowers with which they have poor morphological trait matching, leading to little or no pollen transfer while reducing floral rewards available for other pollinators (Stang et al. 2009). Second, there is the extreme example of nectar robbing, an antagonistic interaction from pollinators wherein the visitors do not visit the flower “legitimately” but rather pierce holes in a flower’s corolla (or utilize a hole that has already been made) to access the nectar rewards without ever touching the reproductive parts of the flower and therefore not acting as a pollinator (Bronstein 2001, Genini et al. 2010, Irwin et al. 2010). On the other side of the interaction, some plants produce chemicals in pollen or nectar that that can be harmful to the bees that visit their flowers, or particularly the development of their offspring, reducing bee fitness (Praz et al. 2008, Sedivy et al. 2011, Haider et al. 2012). Third, the benefit that pollinators have on plant reproduction is sensitive to how “faithful” pollinators are to particular plant species in a single foraging bout. Most pollinators are generalist foragers that can, in some contexts, switch between plant species within a single foraging bout (Waser et al. 1996, Brosi & Briggs 2013, Brosi 2016). When pollinators are promiscuous within a single foraging bout, they may transfer heterospecific pollen to floral stigmas, which can have negative effects on both male and female elements of plant reproduction (Morales & Traveset 2008, Flanagan et al. 2011, Arceo-Gómez & Ashman 2011, Brosi & Briggs 2013). While heterospecific pollen deposition is highly variable in nature (Ashman & Arceo-Gomez 2013, Briggs et al. 2016), it can represent a substantial percentage of total pollen on a stigma, often more than 50% of grains (Ashman & Arceo-Gomez 2013).

It has long been recognized that exploitation of mutualisms (“cheating”) is commonplace and can have substantial impacts on the evolutionary persistence of mutualisms (Bronstein 1994, Richardson 2004, Bronstein et al. 2006, Chamberlain et al. 2014). While our understanding of the extent to which antagonistic interactions between plants and their pollinators is not complete, the examples listed above are common enough that the inclusion of such demonstrated negative interactions on network dynamics and how they might impact the robustness of interactions to extinctions warrants exploration.

To our knowledge, there are only two network simulation studies that incorporate the possibility for antagonistic interactions between plants and pollinators (Campbell et al. 2012, Montesinos-Navarro et al. 2017). Of these, Campbell et al. (2012) focused on network assembly and did not consider robustness. While Montesinos-Navarro et al. (2017) did include robustness as an outcome, they focused on plant-parrot interactions (with relatively modest species richness), with a starting assumption of negative interactions altered to include the potential for benefits to plants including pollination and also seed dispersal. Our study adds to those by incorporating antagonistic interactions into multiple empirical plant-pollinator networks consisting of many plant and pollinator species and in which the starting assumption of interaction direction is positive. We build on binary network simulation modeling approaches to assess how the addition of realistic antagonistic interactions in plant-pollinator networks can impact the effects of pollinator species losses on plant species persistence, i.e. network robustness. In previous simulations where all network interactions are considered positive, the removal of a given pollinator species could result only in the loss of one or more plants. By contrast, after incorporating antagonistic interactions, extinction cascades, (i.e. a second round of pollinator extinctions) also become possible (though see Vieira & Almeida-Neto 2015).

Our study examines the overall robustness of plant-pollinator interactions to extinctions in two ways (1) the area under each extinction curve, designated as *R* (i.e. the rate of decline in plant species richness as pollinator species are lost) (Dunne & Williams 2009); and (2) extinction cascade length (i.e. higher order extinctions that occur beyond the induced pollinator knockout and the resulting plant extinction).

We examined the effects of two factors on network robustness: (1) the proportion of negative interactions in the network, including an all-interactions-positive control; and (2) the order of extinction, random pollinator losses vs. specialist-to-generalist vs. generalist-to-specialist. Removing specialists first could be the most probable extinction sequence (Dunne et al. 2002) as specialist pollinators also tend to be the rarest species (e.g. Vazquez & Aizen 2003). By contrast, generalists are thought to be the “backbone” of networks and when highly connected nodes are lost, networks are expected to collapse rather quickly (Dunne et al. 2002, Tylianakis et al. 2010, Albert et al. 2013); but see (Aizen et al. 2012) who show that loss of specialists can *also* accelerate the rate of species loss overall. While losing generalist pollinators first from a network may seem unlikely, there have been rapid declines and range contractions (leading to local extinctions) in several highly generalist bumble bee species which had previously been abundant (Goulson et al. 2008, Meeus et al. 2011).

We hypothesized that first, increasing the number of antagonistic interactions in the network would lead to both a decrease in the robustness of the network (*R,* or the area under the extinction curve) and greater number of extinction cascades. Second, we hypothesized that inclusion of antagonistic interactions would not change the effects of extinction order (specialist-to-generalist, generalist-to-specialist, or random) relative to networks that assumed strictly beneficial interactions (Memmott et al. 2004, 2007).

## Methods

### Empirical networks

Following previous binary network assessments of robustness (Memmott et al. 2004, 2007), we used empirical network datasets to conduct our robustness assessments. We selected three plant-pollinator networks of varying size and connectance that represent a range of natural plant-pollinator interactions; as in previous assessments, this selection is not meant to be exhaustive (Memmott et al. 2004, 2007, Valdovinos et al. 2012). First, the Clemens and Long (1923) network was collected on Pikes Peak, Rocky Mountains, Colorado USA. This is the largest network with 97 plant species forming 918 unique pairwise interactions with 275 pollinator species. Data were collected in various subalpine habitats at 2500 m elevation over 11 years (Clements & Long 1923). Second, the Arroyo et al (1982) network data were collected at an elevation between 2200m and 2600m between 1980 and 1981 in the alpine (Andean) zone of Cordon del Cepo in Central Chile. The network is intermediate in size with 87 plant species forming 372 unique pairwise interactions with 98 pollinator species. Third, the Dupont network (2003) is the smallest, and data were collected in the sub-alpine desert above 2000 m on the island of Tenerife, Canary Islands between May 7 and June 7, 2001. The network consists of 11 plant species forming 109 unique pairwise interactions with 38 pollinator species. These network data sets were retrieved from the NCEAS Interaction Web Database (http://www.nceas.ucsb.edu/interactionweb, September 1, 2013).

### Assignment of antagonistic interactions

To simulate antagonistic interactions between plants and pollinators, we randomly assigned negative values to the existing interactions in each binary empirical network (non-existing interactions were not subject to negative assignment). Negative interactions were set at a value of −1, i.e. equivalent magnitude to positive interactions. Thus, each possible interaction in a network could have a value of −1, 0, or +1. We assumed that the presence and sign of interactions were symmetric between plants and pollinators following network literature that assumes quantitative interaction strengths are symmetric (e.g. Okuyama & Holland 2008, Holland & Hastings 2008). We assessed four different proportions of negative interactions: 0 (control), 0.05, 0.10, and 0.15, with 50 replicate configurations of randomly-assigned negative values for each network at each proportion of negative values. We thus used a total of 3 (empirical networks) × 4 (proportion negative interactions) × 50 (replicate configurations) = 600 networks. We only used configurations of negative interactions that were initially stable, i.e. which would not automatically lead to extinctions in the absence of perturbations. As a stability condition throughout our simulations, we assumed that plants and pollinators need a positive balance of interactions, that is, a row or column sum of greater than or equal to one. By definition, this automatically excluded all highly specialist nodes (i.e. those two or fewer interactions) from being assigned a negative interaction. While the proportion of negative interactions in empirical networks has not been studied to our knowledge, the maximum value we used (15% negative) represents a practical upper limit of negative interaction assignment that allows for stable initial network configurations without excessive search times.

### Extinction simulations

We simulated extinctions by sequentially removing pollinator species one at a time (i.e. pollinator “knockouts”) and recording the number of plant species that were left with a positive sum of pollinator interactions. Plant species left with an interaction sum less than or equal to zero were then considered extinct and removed from the network due to assumed failure to sexually reproduce. Next, we evaluated if the secondary removal of those plant species left a pollinator species with an interaction sum less than or equal to zero. If yes, they were then considered extinct and removed from the network. These cycles of extinctions are referred to throughout the text as “extinction cascades”. This cycle continued until all plant and pollinator species were left with an interaction sum greater than 0 at which point the simulation moved on to the next pollinator species knockout. Knockouts were repeated until all of the plant species were lost from the network. We carried out the extinction simulations separately for each of the aforementioned 600 network configurations and each of the three extinction orders.

### Analysis of network robustness metrics

We evaluated network robustness via two response variables: (1) *R* (the area under the curve of extinction) – this is a quantitative measure of robustness of a network following a species knockout (extinction). (2) Extinction cascades – the number of higher order extinction cycles that take place after a single pollinator species knockout. We examined how both robustness and extinction cascade length were affected by: 1) increasing the proportion of antagonistic interactions; and 2) extinction order (generalists first, specialists first, random); as well as 3) the interaction between these two factors.

#### R: area under the extinction curve

*R* is the sum of the remaining plant species at each time step along the extinction simulation, standardized by its theoretical maximum value, the starting number of plants × the starting number of pollinators (Burgos et al. 2007). We calculated *R* for each of the 50 simulations per order and proportion of antagonistic interactions for all three networks. We used binomial GLMs with a logit link function to statistically assess network robustness.

#### Total number of extinction cascades

Cascade length is based on the higher order extinctions that occurred beyond the induced pollinator knockout and the resulting plant extinction(s). We define cascade length as the extent to which a knockout impacted trophic levels (plant or pollinator) beyond the immediately impacted one. Cascades of length 0 (i.e., no cascade) indicate only direct plant extinctions driven by a given pollinator knockout. A cascade of length 1 results when a pollinator knockout generates plant extinctions which then lead directly to subsequent pollinator extinction(s), while a cascade of length 2 indicates additional subsequent plant extinction(s). A length 3 cascade would continue with additional pollinator extinctions driven by the plant extinctions in the length 2 cascade, and so on. We underscore that to be considered part of an extinction cascade, extinctions must have resulted from a single pollinator knockout. We calculated the total number of cascade events that occurred across the entire knockout sequence for each of the 50 replicate network configurations for each set of starting networks using GLMs with Poisson errors and a log link function.

## Results

First, we examined how the inclusion of antagonistic interactions impacted network robustness (R) and extinction cascades. For all three of the empirical networks, the incorporation of antagonistic interactions decreases *R* (i.e the decline in remaining plant species accelerates as pollinator species are removed) compared to networks that only include mutualistic interactions (Proportion of Antagonistic Interactions; *p* < 10^-5^, models R1-R3, Table 1, Fig 1).

**Table 1:** GLM results for network robustness (*R*) for each network separately.

| Model Number | Network | Variable | Estimate | Std. Error | Z value | P value |
| --- | --- | --- | --- | --- | --- | --- |
| R1 | Dupont | Intercept | 0.999 | 0.013 | 77.21 | $p < 10^{-5}$ |
| | | Proportion Antagonistic | -1.76 | 0.136 | -12.958 | $p < 10^{-5}$ |
| | | Generalist First | -0.637 | 0.017 | -36.48 | $p < 10^{-5}$ |
| | | Specialist First | 0.822 | 0.021 | 39.246 | $p < 10^{-5}$ |
|  |  | Proportion Antagonistic*Generalist First | -0.153 | 0.184 | -0.829 | 0.47 |
|  |  | Proportion Antagonistic*Specialist First | -0.088 | 0.218 | -0.404 | 0.686 |
| R2 | Clemens | Intercept | 0.519 | 0.001 | 347.052 | $p < 10^{-5}$ |
| | | Proportion Antagonistic | -5.333 | 0.016 | -335.789 | $p < 10^{-5}$ |
| | | Generalist First | -1.359 | 0.002 | -619.182 | $p < 10^{-5}$ |
| | | Specialist First | 1.309 | 0.002 | 531.221 | $p < 10^{-5}$ |
| | | Proportion Antagonistic*Generalist First | 2.564 | 0.024 | 107.368 | $p < 10^{-5}$ |
| | | Proportion Antagonistic*Specialist First | -4.988 | 0.025 | -202.52 | $p < 10^{-5}$ |
| R3 | Arroyo | Intercept | 0.665 | 0.003 | 248.79 | $p < 10^{-5}$ |
| | | Proportion Antagonistic | -5.099 | 0.028 | -181.276 | $p < 10^{-5}$ |
| | | Generalist First | -1.31 | 0.004 | -343.465 | $p < 10^{-5}$ |
| | | Specialist First | 1.122 | 0.004 | 257.603 | $p < 10^{-5}$ |
| | | Proportion Antagonistic*Generalist First | 2.876 | 0.041 | 70.217 | $p < 10^{-5}$ |
| | | Proportion Antagonistic*Specialist First | -3.436 | 0.044 | -78.825 | $p < 10^{-5}$ |

**Figure 1:**
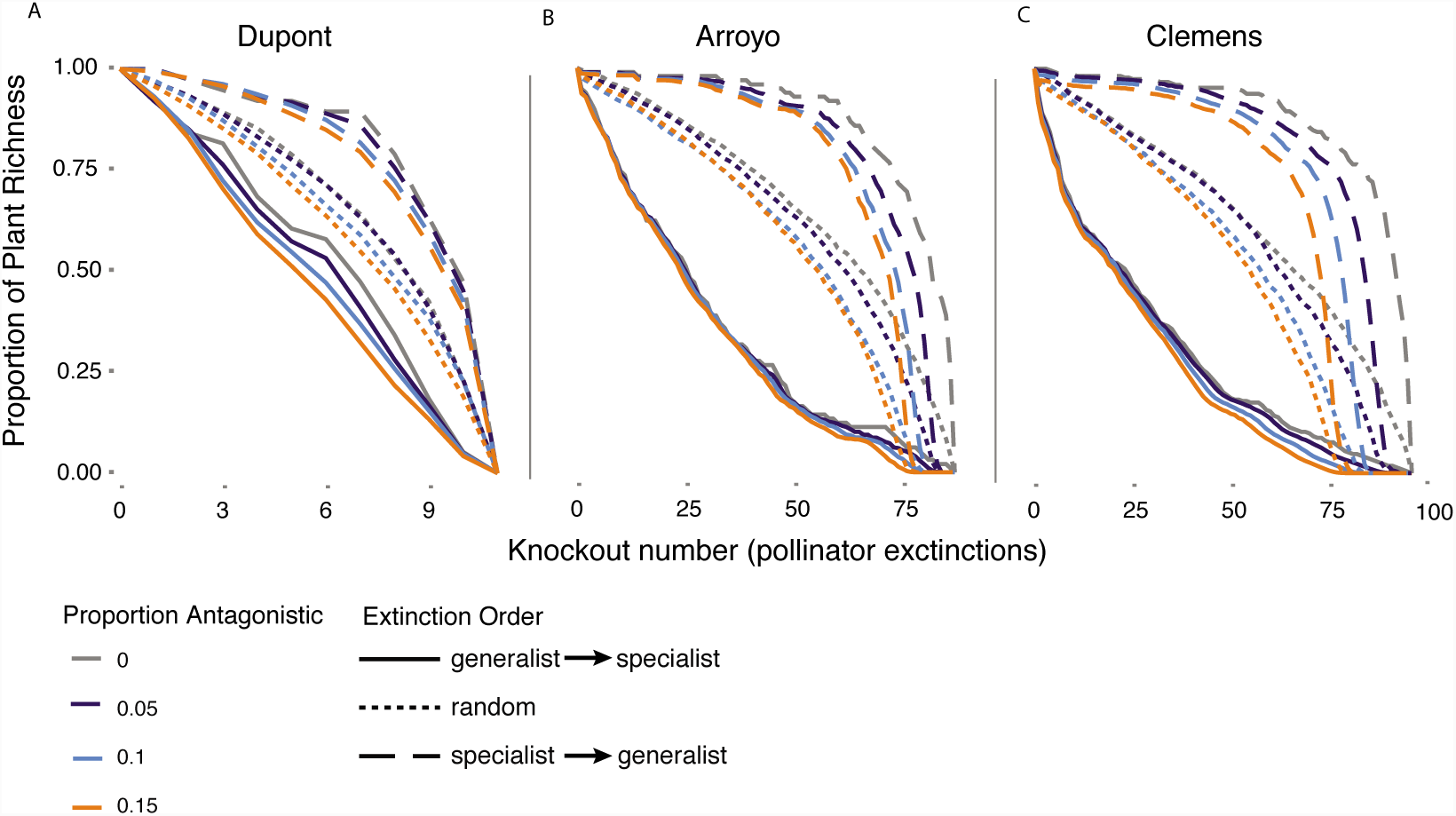
Extinction patterns for the three pollination networks (a) Dupont, (b) Arroyo and (c) Clemens. For a given treatment, we tracked the dynamics of plant extinctions following simulated pollinator knockouts. X axis for each network is constrained by the total number of pollinators in that given network. Lines represent the mean of 50 simulations for each given proportion negative and extinction order.

The effect of antagonistic interactions on extinction cascades differed between networks. In both the Arroyo and Dupont networks, the addition of antagonistic interactions increased the magnitude of extinction cascades (*p* < 10^-5^, Table 1, Fig 2) when compared to those networks that were comprised of all mutualistic interactions. In contrast, the addition of antagonistic interactions had no effect on the number of extinction cascades in the Clemens network (p > 0.5, Table 1, Fig 2).

**Figure 2.**
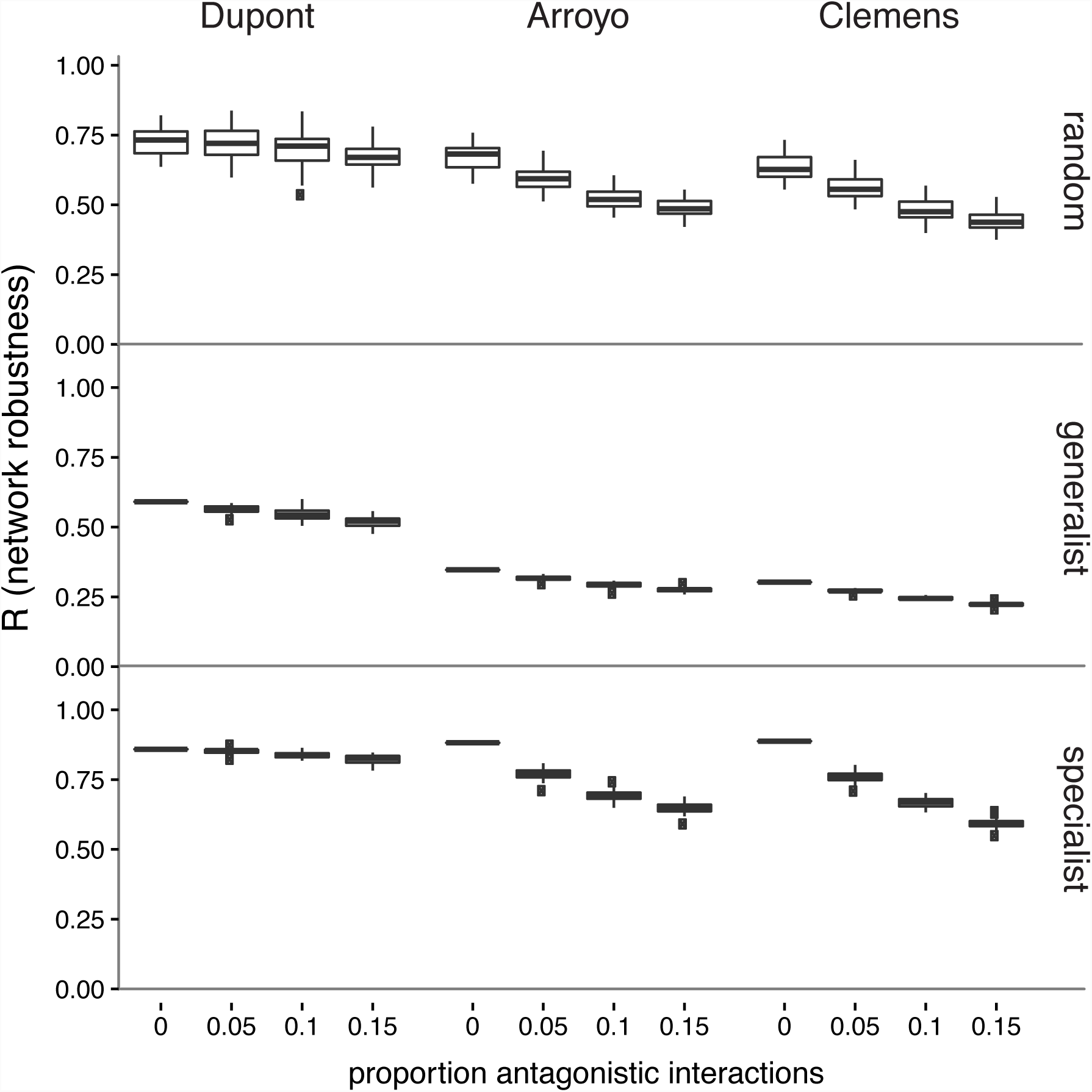
Effects of negative interactions on network robustness during pollinator knockout simulations (shown Figure 1) for different extinction orders and starting networks. Data shown are boxplots displaying median, 50%, and 95% quantiles of robustness (*R*) for the 50 simulations for each proportion negative across the three simulated extinction orders (row headers) and starting network IDs (column headers). Outliers (data points beyond ± 95% quantiles) are displayed as points.

**Figure 3.**
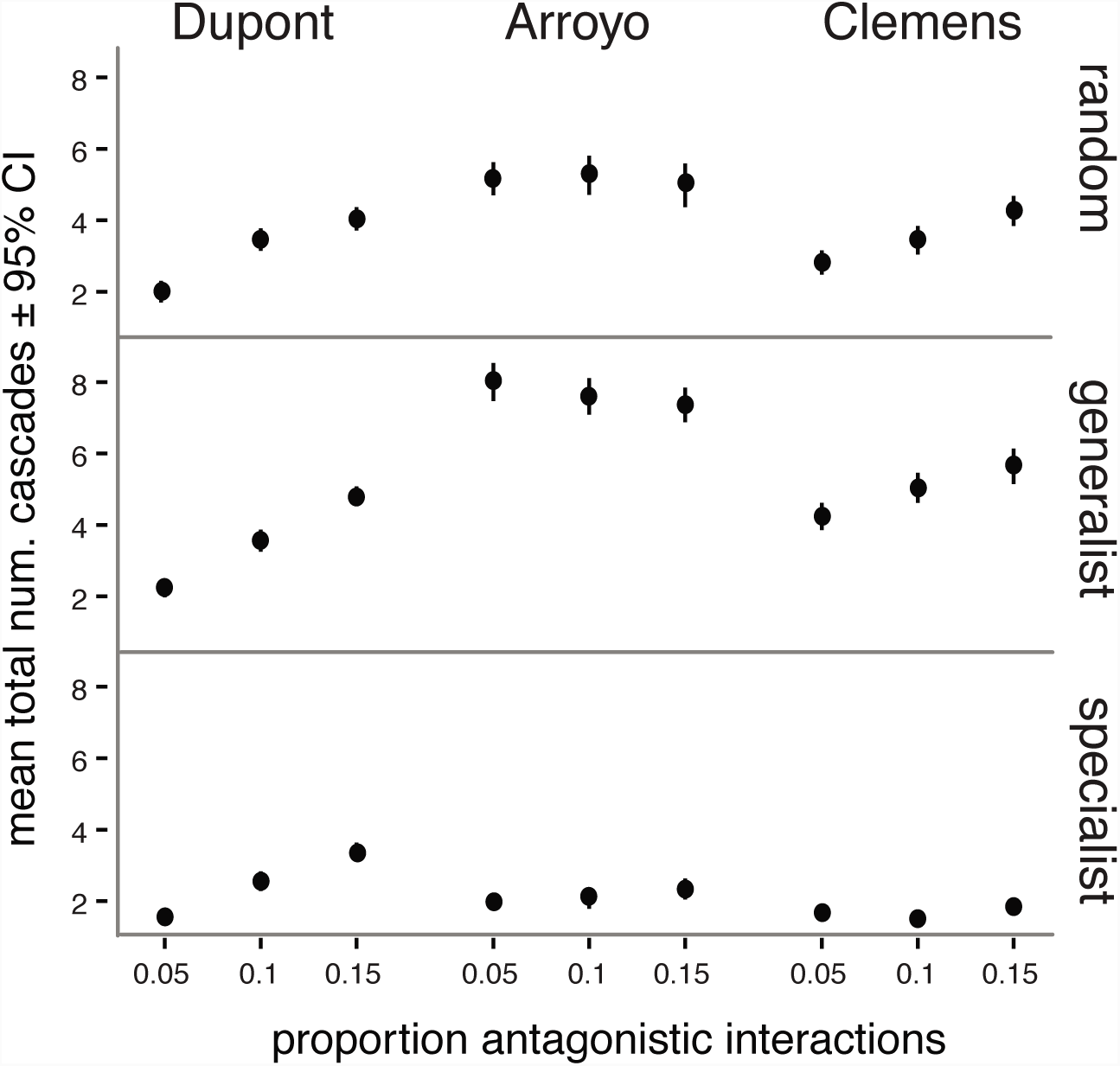
Effects of negative interactions on total number of extinction cascades occurring during pollinator knockout simulations for different extinction orders and starting networks. Points indicate the mean total number of cascades summed across an entire knockout sequence for 50 extinction simulations. Error bars indicate ± 95% confidence intervals. Total cascades are presented for each proportion negative interaction across three simulated extinction orders (row headers) and starting network IDs (column headers).

Next, we examined how extinction order impacted network robustness (*R*) and extinction cascades. When compared to random extinction order, simulations with generalists removed first have the strongest negative impact on *R* (i.e. the intercepts for *R* are lower; p < 10^-5^, Table 1), while removing specialists first has the least negative impact on network robustness (p < 10^-5^, Table 1). This pattern held across all three networks (models R1-R3, Table 1).

The three networks varied in the degree to which extinction order impacted the number of extinction cascades produced after knockouts. For the Clemens network, when the generalists were removed first, the total number of extinction cascades was higher than in the other two orders (main effect *p* < 10^-5^, model C2, Table 2). In contrast, when specialists were removed first we saw the fewest extinction cascades (main effect *p* < 10^-5^, model C2, Table 2). We saw similar results for the Arroyo network; when the generalists were removed first, the total number of extinction cascades was higher than in the other two orders (main effect *p* = .002, model C3, Table 2). When specialists were removed first we saw fewer extinction cascades (main effect *p* = 0.049, model C2, Table 2). Finally, for the Dupont network, extinction order did not impact the total number of extinction cascades (order main effects, *p* > 0.1, model C1, Table 2).

**Table 2.**
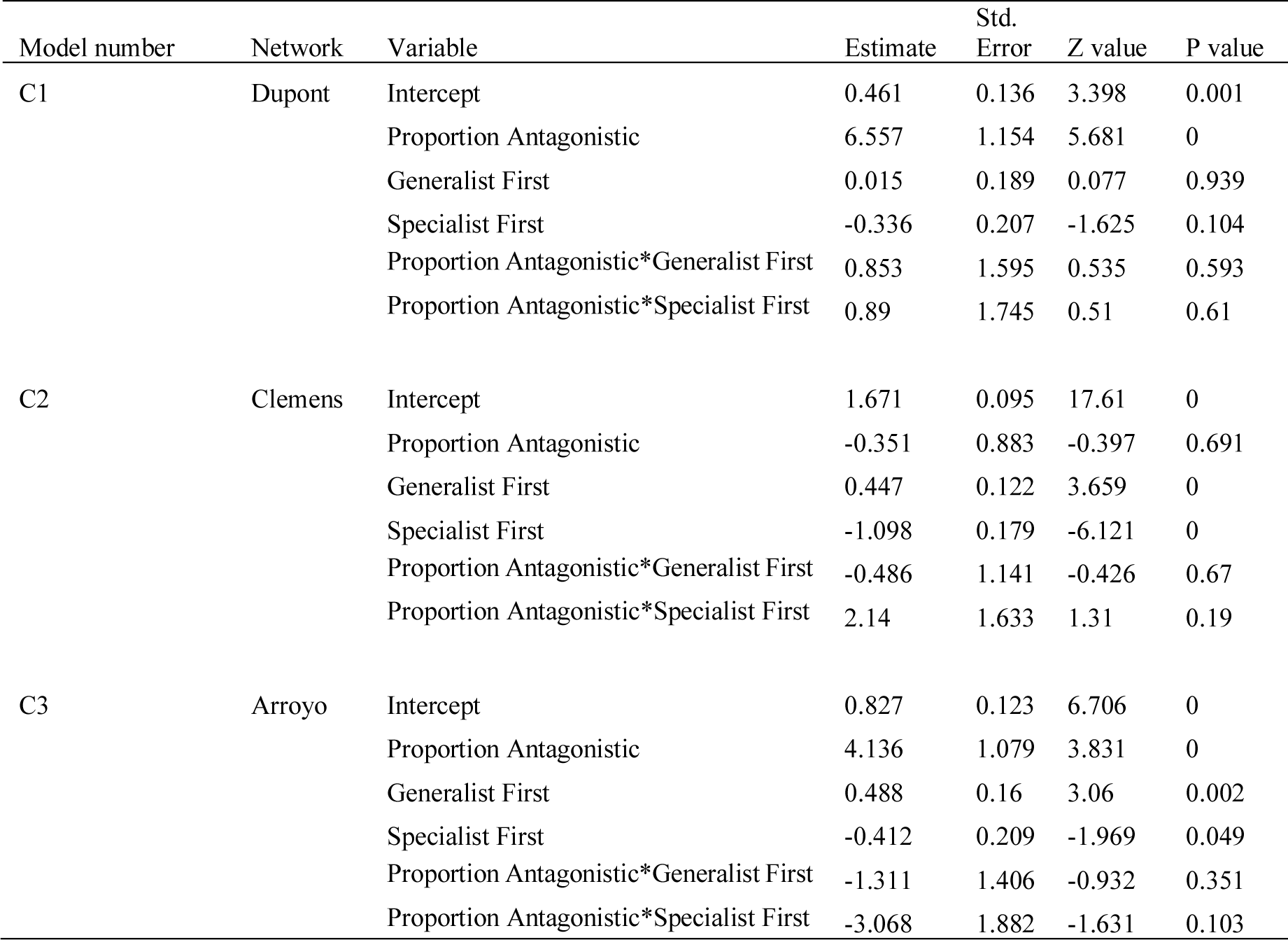
GLM results for total number of cascades vs. proportion antagonistic interactions for each network separately.

| Model number | Network | Variable | Estimate | Std.<br>Error | Z value | P value |
| --- | --- | --- | --- | --- | --- | --- |
| C1 | Dupont | Intercept | 0.461 | 0.136 | 3.398 | 0.001 |
|  |  | Proportion Antagonistic | 6.557 | 1.154 | 5.681 | 0 |
|  |  | Generalist First | 0.015 | 0.189 | 0.077 | 0.939 |
|  |  | Specialist First | -0.336 | 0.207 | -1.625 | 0.104 |
|  |  | Proportion Antagonistic*Generalist First | 0.853 | 1.595 | 0.535 | 0.593 |
|  |  | Proportion Antagonistic*Specialist First | 0.89 | 1.745 | 0.51 | 0.61 |
| C2 | Clemens | Intercept | 1.671 | 0.095 | 17.61 | 0 |
|  |  | Proportion Antagonistic | -0.351 | 0.883 | -0.397 | 0.691 |
|  |  | Generalist First | 0.447 | 0.122 | 3.659 | 0 |
|  |  | Specialist First | -1.098 | 0.179 | -6.121 | 0 |
|  |  | Proportion Antagonistic*Generalist First | -0.486 | 1.141 | -0.426 | 0.67 |
|  |  | Proportion Antagonistic*Specialist First | 2.14 | 1.633 | 1.31 | 0.19 |
| C3 | Arroyo | Intercept | 0.827 | 0.123 | 6.706 | 0 |
|  |  | Proportion Antagonistic | 4.136 | 1.079 | 3.831 | 0 |
|  |  | Generalist First | 0.488 | 0.16 | 3.06 | 0.002 |
|  |  | Specialist First | -0.412 | 0.209 | -1.969 | 0.049 |
|  |  | Proportion Antagonistic*Generalist First | -1.311 | 1.406 | -0.932 | 0.351 |
|  |  | Proportion Antagonistic*Specialist First | -3.068 | 1.882 | -1.631 | 0.103 |

Finally, we assessed the interaction between proportion of antagonistic interactions and extinction order in terms of both *R* and extinction cascade length. Interaction effects were idiosyncratic across the three networks in terms of *R*. For the Dupont network, the addition of antagonistic interactions had no effect on *R* across all extinction orders (interaction terms *p* > 0.5). In both the Arroyo and Clemens networks, the addition of antagonistic interactions dramatically reduced *R* when specialists were removed first compared to the random extinction order simulations (interaction terms both *p* < 10^-5^, models R2-3; Table 1, Fig 2). In contrast, in both the Arroyo and Clemens networks, when generalists were removed first, the effect of increasing negative interactions was reduced relative to the random extinction order (interaction terms both *p* < 10^-5^, models R2– 3; Table 1, Fig 2). In contrast, the impact of extinction order on extinction cascades was not affected by the proportion of antagonistic interactions (interaction terms *p* > 0.4, models C1-C3, Table 2, Fig 2). This was true for all three networks.

## Discussion

Our study examined how both the incorporation of antagonistic interactions into networks, as well as order of extinction in extinction simulations, impacted robustness and total number of extinction cascades in three empirical networks. We found that incorporating antagonistic interactions leads to lower network robustness (*R*), i.e. an increased rate of plant species loss (as compared to networks with only mutualistic interactions) in all three of the networks. Furthermore, when compared to random extinction order, simulations with generalists removed first show the lowest overall robustness whereas the removal of specialists first has the least impact on lowering network robustness. This is true for all three networks and followed our expectations based on previous network extinction simulations (Memmott et al. 2004, 2007). Finally, the addition of increasingly antagonistic interactions did not always magnify the effects of extinction order, leading to idiosyncratic results across the three networks.

While some of the results from our simulations are intuitive (e.g. addition of antagonistic interactions could make networks less robust to extinction), not all of our results could be predicted *a priori*. We did not expect the networks to behave idiosyncratically with respect to how antagonistic interactions and extinction order impacted both *R* and the total number of extinction cascades. Specifically, we found that in the smallest of the networks (Dupont), extinction order did not impact the magnitude of the effect of antagonistic interactions on network robustness. The effect of removing generalists first from the two larger networks (Arroyo and Clemens) largely overshadowed the impact of antagonistic interactions, leading us to conclude that the impact of losing generalists can, in some cases, be so detrimental so as to make inclusion of antagonistic interactions essentially irrelevant. Interestingly, across the three networks the impact of antagonistic interactions on total cascade length was unpredictable. Neither increasing antagonistic interactions nor extinction order were a good predictor for how many extinction cascades would take place in a simulation. Total number of extinction cascades is likely related to structural properties unique to each network, which warrant further exploration.

In all three networks, removing pollinator species from generalist to specialist first has a larger impact on the robustness of the network than with the order of specialist to generalist removals. This pattern has been noted in previous studies (Memmott et al. 2004 and 2007) though ours is the first study to examine the role of extinction order after incorporating antagonistic interactions in the networks. Memmott et al. (2004 and 2007) noted from their study (again, which included only positive interactions) that while robustness was impacted when species were removed from generalist to specialist, the effect was not as dramatic as expected or as reported in food web studies when the most linked interactors are removed (Dunne et al. 2002, Curtsdotter et al. 2011). In our case, adding antagonistic interactions in to the network made the network less robust, resulting in patterns more like those of food web studies where removal of the most linked species causes a collapse to low richness (Dunne et al. 2002, Srinivasan et al 2007).

To our knowledge, there are only two network studies that incorporate antagonistic interactions into network simulations (Campbell et al. 2012, Montesinos-Navarro et al. 2017). In their models, Campbell et al. (2012) define interactions as either mutually beneficial or beneficial for one species and detrimental to the other (in contrast to our model, which compares the robustness of networks in which all interactions are considered beneficial to those that incorporate the possibility of antagonistic interactions). Another key difference between our work and the Campbell model is their focus on hypothetical (rather than empirical) networks. While their model is focused on network assembly, not robustness to co-extinctions, their simulations revealed “critical species”—those species that cause significant community collapse when removed—as species that tend to have asymmetric interaction direction. Their results suggest that when these critical species are lost, the network is left with an abundance of antagonistic interactions in their absence, which can lead to further extinctions and often collapse. In the second study, Montesinos-Navarro et al. (2017) assessed the relative contribution of antagonistic vs mutualistic interactions to nestedness and modularity in plant-parrot interactions, which were first assumed to be negative, with positive interactions (seed dispersal, pollination) then added. Through simulated coextinction cascades, Montesinos-Navarro et al. revealed that in simulations where the beneficial effects of parrots were considered, the networks were more robust to extinctions. These findings, considered in conjunction with the results presented here, underscore the importance of considering the range of interactions types (mutualist to antagonistic) that can be realized in plants—animal interactions.

To further our understanding of how the incorporation of antagonistic interactions could affect robustness to co-extinction, we suggest two lines of inquiry for future studies: 1) explicit consideration of network structural properties; and 2) incorporation of re-wiring dynamics. In terms of the first area, structural attributes including nestedness, degree distribution, connectance, and many others can drive network outcomes such as persistence (e.g. Valdovinos et al. 2016), as well as local stability and resilience (e.g. Okuyama and Holland 2008). While our work considered three distinct plant-pollinator networks—with contrasting levels of connectance, species richness, and other attributes—it was beyond the scope of this study to directly consider the effects of network structure. Future work should address how different network structural properties operate in conjunction with antagonistic interactions to shape robustness. Second, a substantial body of work has revealed that re-wiring (potential for change in interaction identity, typically following partner extinctions or substantial changes in abundance) can alter network robustness to co-extinctions (Fortuna & Bascompte 2006, Kaiser-Bunbury et al. 2010). Again, while inclusion of re-wiring dynamics was beyond the scope of this study, its consideration could change our understanding of how antagonistic interactions impact network robustness. In particular, given that re-wiring tends to buffer networks against coextinctions, it could help to offset the negative effects of antagonistic interactions.

Habitat loss and degradation are leading to global pollinator losses (Biesmeijer et al. 2006; Potts et al.2010). Given the central importance of pollination in food production and the maintenance of biodiversity and ecosystem function, it is imperative that we come to a predictive under-standing of how pollinator losses will affect plant-pollinator systems. Evidence suggests that losing even a few pollinators can potentially have a strong antagonistic effect on the plants that rely on pollination for reproduction (Biesmeijer et al. 2006, Potts et al. 2010, Brosi & Briggs 2013). In the absence of empirical studies, network-based simulations suggest some potential that plant communities could be robust to pollinator extinctions (Memmott et al. 2004, 2007, Kaiser-Bunbury et al. 2010). While not exhaustive, this study reveals how one key assumption in plant-pollinator simulation models may contribute to overestimation of network robustness. By incorporating more realistic representations of the interactions that take place in a plant-pollinator community, we are more likely to identify properties of networks that determine robustness to extinctions. Future studies that improve on predictive models will allow us to anticipate likely changes in pollination services and help us design strategies to maximize ecosystem resilience.

## Acknowledgements

This work was assisted by attendance as a Short-term Visitor at the National Institute for Mathematical and Biological Synthesis, an Institute supported by the National Science Foundation through NSF Award #DBI-1300426, with additional support from The University of Tennessee, Knoxville. In addition, this work was supported by a Doctoral Fellowship to HMB from the Institute of Quantitative Theory and Methods at Emory University. P. Humphrey provided invaluable feedback on the models and manuscript.

## References

Aizen, M. A., M. Sabatino, and J. M. Tylianakis. 2012. Specialization and rarity predict nonrandom loss of interactions from mutualist networks. Science 335:1486–1489.

Albert, E. M., M. A. Fortuna, J. A. Godoy, and J. Bascompte. 2013. Assessing the robustness of networks of spatial genetic variation. Ecology Letters 16:86–93.

Arceo-Gómez, G., and T.-L. Ashman. 2011. Heterospecific pollen deposition: does diversity alter the consequences? New Phytologist 192:738–746.

Ashman, T. L., and G. Arceo-Gomez. 2013. Toward a predictive understanding of the fitness costs of heterospecific pollen receipt and its importance in co-flowering communities. American Journal of Botany 100:1061–1070.

Bascompte, J., and P. Jordano. 2007. Plant-animal mutualistic networks: The architecture of biodiversity. Annual Review of Ecology Evolution and Systematics 38:567–593.

Bascompte, J., P. Jordano, and J. Olesen. 2006. Asymmetric coevolutionary networks facilitate biodiversity maintenance. Science 312:431–433.

Bascompte, J., P. Jordano, C. J. Melián, and J. M. Olesen. 2003. The nested assembly of plant-animal mutualistic networks. Proceedings of the National Academy of Sciences 100:9383–9387.

Biesmeijer, J. C., S. P. M. Roberts, M. Reemer, R. Ohlemueller, M. Edwards, T. Peeters, A. P. Schaffers, S. G. Potts, R. Kleukers, C. D. Thomas, J. Settele, and W. E. Kunin. 2006. Parallel declines in pollinators and insect-pollinated plants in Britain and the Netherlands. Science 313:351–354.

Briggs, H. M., L. M. Anderson, L. M. Atalla, A. M. Delva, E. K. Dobbs, and B. J. Brosi. 2016. Heterospecific pollen deposition in Delphinium barbeyi: linking stigmatic pollen loads to reproductive output in the field. Annals of Botany 117:341–347.

Bronstein, J. L. 1994. Conditional outcomes in mutualistic interactions. Trends in Ecology & Evolution 9:214–217.

Bronstein, J. L. 2001. The exploitation of mutualisms. Ecology Letters 4:227–287.

Bronstein, J. L., R. Alarcón, and M. Geber. 2006. The evolution of plant–insect mutualisms. New Phytologist 172:412–428.

Brosi, B. J. 2016. Pollinator specialization: from the individual to the community. New Phytologist 210:1190–1194.

Brosi, B. J., and H. M. Briggs. 2013. Single pollinator species losses reduce floral fidelity and plant reproductive function. Proceedings of the National Academy of Sciences 110:13044–13048.

Burgos, E., H. Ceva, R. P. J. Perazzo, M. Devoto, D. Medan, M. Zimmermann, and A. María Delbue. 2007. Why nestedness in mutualistic networks? Journal of Theoretical Biology 249:307–313.

Campbell, C., S. Yang, K. Shea, and R. Albert. 2012. Topology of plant-pollinator networks that are vulnerable to collapse from species extinction. Physical Review E 86:021924.

Chamberlain, S. A., J. L. Bronstein, and J. A. Rudgers. 2014. How context dependent are species interactions? Ecol Lett 17:881–890.

Dunne, J. A., and R. J. Williams. 2009. Cascading extinctions and community collapse in model food webs. Philosophical Transactions of The Royal Society B-Biological Sciences 364:1711–1723.

Dunne, J. A., R. J. Williams, and N. D. Martinez. 2002. Food-web structure and network theory: The role of connectance and size. Proceedings of the National Academy of Sciences 99:12917–12922.

Flanagan, R. J., R. J. Mitchell, and J. D. Karron. 2011. Effects of multiple competitors for pollination on bumblebee foraging patterns and Mimulus ringens reproductive success. Oikos 120:200–207.

Fortuna, M., and J. Bascompte. 2006. Habitat loss and the structure of plant-animal mutualistic networks. Ecology Letters 9:278–283.

Genini, J., L. P. C. Morellato, P. R. Guimarães, and J. M. Olesen. 2010. Cheaters in mutualism networks. Biology Letters 6:494–497.

Goulson, D., G. C. Lye, and B. Darvill. 2008. Decline and conservation of bumble bees. Annual Review of Entomology 53:191–208.

Holland, M. D., and A. Hastings. 2008. Strong effect of dispersal network structure on ecological dynamics. Nature 456:792–794.

Irwin, R. E., J. L. Bronstein, J. S. Manson, and L. Richardson. 2010. Nectar Robbing: Ecological and Evolutionary Perspectives. Annual Review of Ecology Evolution And Systematics 41:271–292.

Kaiser-Bunbury, C. N., S. Muff, J. Memmott, C. B. Müller, and A. Caflisch. 2010. The robustness of pollination networks to the loss of species and interactions: a quantitative approach incorporating pollinator behaviour. Ecology Letters 13:442–452.

Meeus, I., M. J. F. Brown, D. C. De Graaf, and G. Smagghe. 2011. Effects of invasive parasites on bumble bee declines. Conservation Biology 25:662–671.

Memmott, J., N. M. Waser, and M. V. Price. 2004. Tolerance of pollination networks to species extinctions. Proceedings Biological sciences / The Royal Society 271:2605–2611.

Memmott, J., P. G. Craze, N. M. Waser, and M. V. Price. 2007. Global warming and the disruption of plant-pollinator interactions. Ecology Letters 10:710–717.

Montesinos-Navarro, A., F. Hiraldo, J. L. Tella, and G. Blanco. 2017. Network structure embracing mutualism-antagonism continuums increases community robustness. Nature Ecology & Evolution 1:1661–1669.

Morales, C., and A. Traveset. 2008. Interspecific Pollen Transfer: Magnitude, Prevalence and Consequences for Plant Fitness. Critical Reviews in Plant Sciences 27:221–238.

Okuyama, T., and J. N. Holland. 2008. Network structural properties mediate the stability of mutualistic communities. Ecology Letters 11:208–216.

Potts, S. G., J. C. Biesmeijer, C. Kremen, P. Neumann, O. Schweiger, and W. E. Kunin. 2010. Global pollinator declines: trends, impacts and drivers. Trends in Ecology & Evolution 25:345–353.

Ramos-Jiliberto, R., F. S. Valdovinos, P. Moisset de Espanés, and J. D. Flores. 2012. Topological plasticity increases robustness of mutualistic networks. Journal of Animal Ecology 81:896–904.

Richardson, S. C. 2004. Are nectar-robbers mutualists or antagonists? - Springer. Oecologia.

Stang, M., P. G. L. Klinkhamer, N. M. Waser, I. Stang, and E. van der Meijden. 2009. Size-specific interaction patterns and size matching in a plant-pollinator interaction web. Annals of Botany 103:1459–1469.

Thierry, A., A. P. Beckerman, P. H. Warren, R. J. Williams, A. J. Cole, and O. L. Petchey. 2011. Adaptive foraging and the re-wiring of size-structured food webs following extinctions. Basic and Applied Ecology 12:562–570.

Tylianakis, J. M., E. Laliberté, A. Nielsen, and J. Bascompte. 2010. Conservation of species interaction networks. Biological Conservation 143:2270–2279.

Valdovinos, F. S., P. Moisset de Espanés, J. D. Flores, and R. Ramos-Jiliberto. 2012. Adaptive foraging allows the maintenance of biodiversity of pollination networks. Oikos 122:907–917.

Valdovinos, F. S., B. J. Brosi, H. M. Briggs, P. Moisset de Espanés, R. Ramos-Jiliberto, and N. D. Martinez. 2016. Niche partitioning due to adaptive foraging reverses effects of nestedness and connectance on pollination network stability. Ecology Letters 19:1277–1286.

Vazquez, D. P., and M. A. Aizen. 2003. Null model analyses of specialization in plant-pollinator interactions. Ecology 9:2493–2501

Waser, N. M., L. Chittka, M. Price, Williams, and J. Ollerton. 1996. Generalization in pollination systems, and why it matters. Ecology 77:1043–1060.

